# Human and mouse regenerative macrophages enhance beta cell survival, function, and proliferation

**DOI:** 10.1101/2025.08.04.668458

**Authors:** Mahdis Monajemi, Alexandra S. Craciun, Qing Huang, Lei Dai, Majid Mojibian, Sarah Q Crome, C. Bruce Verchere, Megan K. Levings

## Abstract

**Aims/hypothesis:** Type 1 diabetes is an autoimmune disease caused by immune-mediated destruction of insulin-producing pancreatic beta cells. Interestingly, individuals with long-standing type 1 diabetes have residual beta cells, suggesting the existence of regenerative mechanisms that help maintain beta cell survival. Islet-resident macrophages have an important role in type 1 diabetes, and during disease progression can adopt a tissue-regenerating phenotype which may support beta cells. However, the specific roles of macrophages in beta cell survival, function, and proliferation remains poorly defined. This study aimed to elucidate how different macrophage subtypes influence beta cell survival, function, and proliferation.

**Methods:** Mouse and human islets were isolated from the pancreas and co-cultured in vitro with macrophages. To investigate whether macrophages enhance beta cell survival and function, beta cell apoptosis was measured using flow cytometry, and insulin secretion was assessed using a glucose-stimulated insulin secretion assay. We also examined whether macrophages further increased beta cell proliferation in the presence of harmine, a DYRK1A inhibitor. Finally, we evaluated the effect of islet co-culture on macrophage phenotype by flow cytometry and cytokine secretion analysis.

**Results:** We found that regenerative, but not pro-inflammatory, macrophages enhanced beta cell survival and function through mechanisms that did not require direct cell contact. Direct contact between macrophages and islets further promoted a regenerative phenotype in macrophages, characterized by increased CD206 expression and secretion of anti-inflammatory factors. Additionally, regenerative macrophages promoted beta cell proliferation in the presence of harmine.

**Conclusions:** Our findings demonstrate that regenerative macrophages support pancreatic beta cell survival, function, and proliferation. Harnessing the regenerative properties of macrophages could offer a novel strategy to promote beta cell survival and function, thereby improving outcomes for individuals with type 1 diabetes.

**Research in Context:** *What is already known about this subject?*

- Macrophages are the predominant resident immune cells within pancreatic islets in non-diabetic individuals; they contribute to tissue homeostasis and immune surveillance.
- Depending on their activation state, macrophages can exert either beneficial (pro-regenerative) or harmful (pro-inflammatory) effects on beta cell function.

*What is the key question?*

- How do different macrophage subtypes regulate islet survival, regeneration, and function?

*What are the new findings?*

- Co-culturing mouse or human islets with regenerative macrophages enhanced beta cell survival and function. When added in the presence of a DYRK1A inhibitor (harmine), they also promoted beta cell proliferation.
- Regenerative macrophage-derived factors promoted mouse islet function via a contact independent mechanism.
- Mouse islets enhanced the regenerative phenotype of macrophages.

*How might this impact on clinical practice in the foreseeable future?*

- Leveraging the regenerative potential of macrophages represents a novel therapeutic approach to enhance beta cells, ultimately improving outcomes for individuals with type 1 diabetes.

## Introduction

Type 1 diabetes is an autoimmune disease characterized by beta cell loss and lifelong insulin dependence [1]. Although T cells are ultimately thought to mediate direct beta cell killing, many different innate and adaptive immune cell subsets are implicated in disease pathogenesis, with therapies to modulate innate and/or adaptive immunity in clinical testing [2]. Interestingly, a consistent finding is that in people with newly diagnosed or long-standing type 1 diabetes, some beta cells remain, raising the possibility that promoting residual beta cell survival, proliferation and/or function could be a therapeutic approach [3, 4] [5].

Macrophages are innate immune cells essential for tissue homeostasis, immunity, and organ development. Found in lymphoid and non-lymphoid tissues, they are highly adaptable cells which adjust their phenotype and function in response to environmental signals [6]. They switch between pro-inflammatory and anti-inflammatory roles as needed, and through the release of cytokines and growth factors and phagocytosis of dead cells, they help regulate inflammation maintain healthy tissues [6–8].

Macrophages are the most abundant immune cells residing within islets of non-diabetic humans [9], and in the NOD mouse model of type 1 diabetes prior to the development of inflammation [10]. Under these steady state conditions, islet-resident macrophages have a pro-inflammatory activated state [11] and respond to signals such as elevated glucose and islet amyloid formation by increasing pro-inflammatory cytokine expression, leading to further beta cell dysfunction and hyperglycemia in rodent models of type 2 diabetes [12] [13–16]. Islet macrophages also play an important role in beta cell loss in the NOD mice [17].

Recent studies show that islet macrophages are not only involved in beta cell death and dysfunction in type 1 diabetes and type 2 diabetes, but also in the regeneration and proliferation of beta cells. Specifically, anti-inflammatory, pro-regenerative macrophages can stimulate tissue regeneration and beta cell proliferation through secretion of molecules such as TGF-β [18]. In obese mice, islet macrophages promote beta cell proliferation through PDGF-PDGFR signaling [19] and VGEF-mediated recruitment of endothelial cells [20]. In mice with streptozotocin-induced diabetes, beta cell death is associated with upregulated IGF-1 expression in islet macrophages, and their depletion leads to impaired glucose tolerance [21]. Also consistent with a regenerative role in diabetes, macrophages mimicking tumor-associated macrophages accelerate wound healing in mouse models of type 1 and 2 diabetes [22]. Mesenchymal stem cells, which also promote pancreatic beta cell regeneration [23], engineered to express human α-1 antitrypsin induce protective macrophages and enhance islet graft survival [24]. Overall, this extensive body of literature supports the concept that macrophages with a pro-regenerative phenotype are beneficial for islet health and function

Thus, depending on the context, macrophages can have either beneficial (pro-regenerative) or detrimental (pro-inflammatory) effects on beta cell regeneration and/or performance. However, the specific roles of different macrophage subtypes in beta cell survival, regeneration, and function remained unclear, as did the role of contact-dependent or - independent mechanisms. To better understand the effects of different macrophage subtypes on beta cell survival and function, we developed an ex vivo co-culture model of macrophages and islet cells. In this study, we show that regenerative macrophages are highly versatile regulators of beta cell function and survival, capable of significantly modulating insulin secretion and islet survival through secreted factors.

## Methods

### Informed consent and ethics committee approval

Human macrophages were generated from buffy coats obtained from Canada Blood Services from healthy volunteers who gave informed consent and with approval from the University of British Columbia Clinical Research Ethics Board (H18-02553). Human islets were collected from the pancreases of deceased organ donors at the IsletCore (University of Alberta) with approval from the University of British Columbia Clinical Research Ethics Board (H20-01786).

### Animals

C57BL/6 mice were sourced from Jackson Laboratory and bred in-house under specific pathogen-free conditions. They were housed in the Animal Research Center at BC Children’s Hospital Research Institute. All experiments were conducted in compliance with institutional guidelines and the standards set by the Canadian Council on Animal Care under protocols A23-0265 and A21-0138.

### Mouse macrophage generation

Macrophages were derived from bone marrow progenitor cells isolated from the femurs and tibias of mice, following previously established methods [25, 26]. To culture the macrophages, bone marrow aspirates were plated at a density of 1× 10⁶ cells/ml in IMDM (Gibco) supplemented with 10% FBS (Gibco), penicillin-streptomycin (Gibco), and 10 ng/ml MCSF (StemCell Technologies). The cells were incubated at 37°C with 5% CO₂ for 10 days. During this period, the media was replaced on days 4 and 7 to maintain optimal growth conditions. Vigorous pipetting with Cell Dissociation Buffer (Gibco) was used to detach adherent BMDMs. Cells were differentiated into M(LPS, IFN-γ) or M(IL-4, IL13) macrophages after incubation with 100 ng/ml of LPS (Sigma-Aldrich), and 20 ng/ml of IFN-γ, IL-4, and IL-13 (all from StemCell Technologies) for 3 days.

### Human macrophage generation

Peripheral blood mononuclear cells (PBMCs) were isolated from blood as described previously [27], monocytes were isolated from PBMCs using the EasySep™ Human CD14 Positive Selection Kit (StemCell Technologies). The monocytes were then plated in RPMI cell culture medium supplemented with 10% FBS, penicillin-streptomycin, and 15mM Hepes (all from Gibco), and 100 ng/ml MCSF (StemCell Technologies) at a density of 150 × 10⁵ cells/mL in an appropriately sized flask and incubated at 37°C for 6 days. On day 7, room temperature Cell Dissociation Buffer (Gibco) was added to the flask to detach the cells. The cells were resuspended at 1.5 x 10^5^ cells/mL and polarized into M(IL-4, IL-13) phenotype after incubation with 20 ng/ml of IL-13 and IL-4 (both from StemCell Technologies) at 37°C for 3 days.

### Apoptosis assay

Apoptosis was assessed using the Apoptosis/Necrosis assay kit (Abcam). Islet cells, dissociated after incubation with Accutase (StemCell Technologies) for 15 min, were stained with Apopxin Green (to identify apoptotic cells) and CytoCalcein Violet 450 (for healthy cells) according to the manufacturer’s instructions, as previously described [3]. Briefly, the dissociated islet cells were stained with Apopxin, CytoCalcein, and CD45 antibody. The staining was performed at room temperature for 30 min. After staining, the cells were washed and acquired using a flow cytometer FACSymphony A5 (BD Biosciences), and data analysis was performed using the FlowJo 10.8.2 (BD Biosciences).

### Proliferation assay

Beta cell proliferation assays were conducted using the Click-iT Plus EdU Flow Cytometry Assay Kit (Invitrogen). Briefly, intact islet cells were incubated with or without macrophages and ±10 µM harmine (Sigma-Aldrich) for 72 hours, and during the final 24 hours incubation, EdU (10 µM) was added. The islets were dissociated with Accutase (StemCell Technologies) for 15 min. Cells were then washed with 1% BSA in PBS and stained with anti-CD45 and fixable viability dye. Cells were fixed with Click-iT fixative for 15 min, followed by a 15 min incubation for permeabilization with Click-iT permeabilizing reagent. Insulin and glucagon staining was performed overnight. Subsequently, cells were washed with permeabilization reagent and the components of the Click iT detection cocktail were added for 30 min. Finally, the cells were washed twice with permeabilization reagent (500 ×g, 5 min) and were resuspended in FACS buffer before acquisition using a flow cytometer FACSymphony A5 (BD Biosciences). Data analysis was performed using the FlowJo 10.8.2 (BD Biosciences).

### Mouse islet isolation and culture

Mice anesthetized with isoflurane were sacrificed by cervical dislocation and islets isolated as previously reported [4]. The common bile duct was clamped at the junction where it meets the intestine, and the pancreas was injected with collagenase XI (Sigma-Aldrich) in HBSS. It was then incubated with collagenase solution at 37°C for 15 min, followed by cold HBSS with calcium chloride to terminate the digestion. The islets were washed, filtered through a 70 µm strainer, and rinsed with islet media to collect them. Finally, the islets were handpicked under a Nikon SMZ800 microscope and transferred to a fresh Petri dish with islet media (RPMI 1640 medium containing 11.1 mM glucose, 2 mM L-Glutamine, Phenol Red) supplemented with 10% FBS, 2 mM L-alanyl-Lglutamine dipeptide (GlutaMAX), 1% penicillin/streptomycin (All from Gibco) at 37°C in 5% CO2. For each experiment, a pool of approximately 100 visually healthy islets (characterized by a round shape, absence of a necrotic core, and consistent brownish color) was used per condition in the in vitro studies. Islets were cultured in islet media containing 5 mM glucose, either alone or co-cultured with macrophages at a ratio of 1 islet to 1000 macrophages.

### Islet C-peptide secretion and content

Islets were isolated via collagenase digestion as previously described. After a 72 hours incubation with or without macrophages, similarly-sized islets per sample were pre-incubated for 60 min in Krebs-Ringer Bicarbonate buffer (KRB) containing 0.1% BSA, pH 7.4, and 1.67 mM glucose at 37°C. Islets were then transferred to fresh KRB containing 1.67 mM glucose for an additional 60 min at 37°C, and the supernatant was collected. For insulin secretion measurements, islets were transferred to 16.7 mM glucose KRB followed by 1 hour incubation with 40 mM KCl KRB. Supernatants were collected at the end of each incubation period. C-peptide content from both the supernatants and islet pellets, extracted using acid ethanol, was measured using mouse C-peptide ELISA (Alpco) and human C-peptide ELISA (Mercodia).

### Multiplex analysis of supernatant cytokines

Mouse and human macrophage cytokines were measured using the LEGENDplex™ multi-analyte bead-based flow assay kit (Biolegend) according to the manufacturer’s instructions. Supernatants were centrifuged to remove any particulates, stored at −20°C, and thawed prior to use. After vortexing the samples and pre-mixed beads, all reagents were prepared at room temperature. Serial 1:4 dilutions of the panel standard were made. Standards and samples were loaded onto a 96-well plate, mixed beads were added, and the plate was incubated in the dark at 600 rpm for 2 hours. After washing, a biotinylated detection antibody was added and incubated for 1 h. Streptavidin–phycoerythrin was added, incubated for 30 min, followed by another wash. The plate was read directly on a flow cytometer FACSymphony A1 (BD Biosciences), and data analysis was performed using the LEGENDplex™ cloud-based software.

### Flow cytometry

Flow cytometry was performed in accordance with the guidelines described by Cossarizza et al. [28]. The antibodies used for flow cytometric analysis are listed in **Table S1**. Briefly, cells were stained for surface markers in the presence of the fixable viability dye eFluor 780 (Thermo Fisher Scientific) to exclude dead cells. Intracellular antigen detection was performed following incubation with a permeabilization buffer (Invitrogen), followed by intracellular staining. Data acquisition was carried out using an LSR Fortessa II, A5 Symphony, or FACSymphony A1 (all from BD Biosciences), and data were analyzed using FlowJo software, version 10.8.2 (BD Biosciences).

### Immunofluorescent staining and imaging

Islets were co-cultured with or without macrophages and ±10 µM harmine (Sigma-Aldrich) in Lab-Tek II Chamber Slide (Sigma-Aldrich) for 72 hours, then washed twice with 3% BSA in PBS. Each chamber was then rinsed twice with 3% BSA in PBS, and fixed in 4% paraformaldehyde (Sigma-Aldrich) for 15 min. After two washes with 3% BSA in PBS, samples were permeabilized for 20 min in a 1% Triton-X 100 solution (Sigma-Aldrich). Each chamber was rinsed twice with 3% BSA in PBS and stained overnight at 4°C with antibodies diluted in PBS (Gibco). Beta and alpha cells were labeled using anti-insulin and anti-glucagon antibodies, respectively, and proliferating cells were labelled using a Ki67 antibody. After staining, samples were washed twice with PBS, and cells were identified using a TO-PRO-3 (Thiazole Red) nuclear stain (Thermo Fisher Scientific). Each incubation was completed with minimal light exposure and at room temperature, unless otherwise specified. After chamber detachment, slides were mounted using Dako mounting medium (Agilent Technologies Canada Inc.) and a 1.5 thickness cover slip. The slides were kept at room temperature for 30 min to allow for solidification of the mounting medium, then stored at 4°C until imaging. The Immunofluorescence antibodies used in the assay are listed **Table S2**. Slides were imaged using the Leica TCS SP8 confocal imaging system, at 25x with water immersion.

### Human islet preparation and culture

Human islets were isolated from the pancreases of deceased organ donors at the IsletCore (University of Alberta) and shipped overnight. Each islet preparation was hand-picked and plated to ensure high purity. Donor characteristics for all individuals used in this study are provided **Table S3**. The islets were cultured in CMRL 1066 medium (Corning) supplemented with 1% Pen/Strep, 1% GlutaMAX, and 0.5% Gentamicin (all from Gibco), with or without human macrophages at a ratio of 1 islet to 1000 macrophages.

### Statistics

Data were analyzed using GraphPad Prism 10.1.1 and are reported as mean ± SEM. Statistical significance was determined using Student’s *t* test (2 tailed), one-way ANOVA, multiple Wilcoxon test, and Friedman test. *P* < 0.05 was considered statistically significant.

## Results

### Mouse M(IL-4, IL13) macrophages improve beta cell proliferation in the presence of harmine

To establish a model of mouse islets co-culture with macrophages of different phenotypes, we first generated mouse bone marrow-derived macrophages (BMDM) which were left unpolarized (M(-)), or polarized to pro-inflammatory (M(LPS, IFN-γ)), or pro-regenerative (M(IL-4, IL-13)) phenotypes, by treatment with LPS and IFN-γ, or IL-4 and IL-13, respectively, for 3 days. Consistent with previous reports [29, 30], M(LPS, IFN-γ) macrophages expressed CD86 and MHC Class II, whereas expression of these markers remained low in M(IL-4, IL-13) and M(-) macrophages **[Fig. 1A, S1A]**. In contrast, M(IL-4, IL-13), but not M(LPS, IFN-γ) or M(-) macrophages, upregulated CD206 **[Fig. 1B, S1A]** [31] [32].

**Figure 1.**
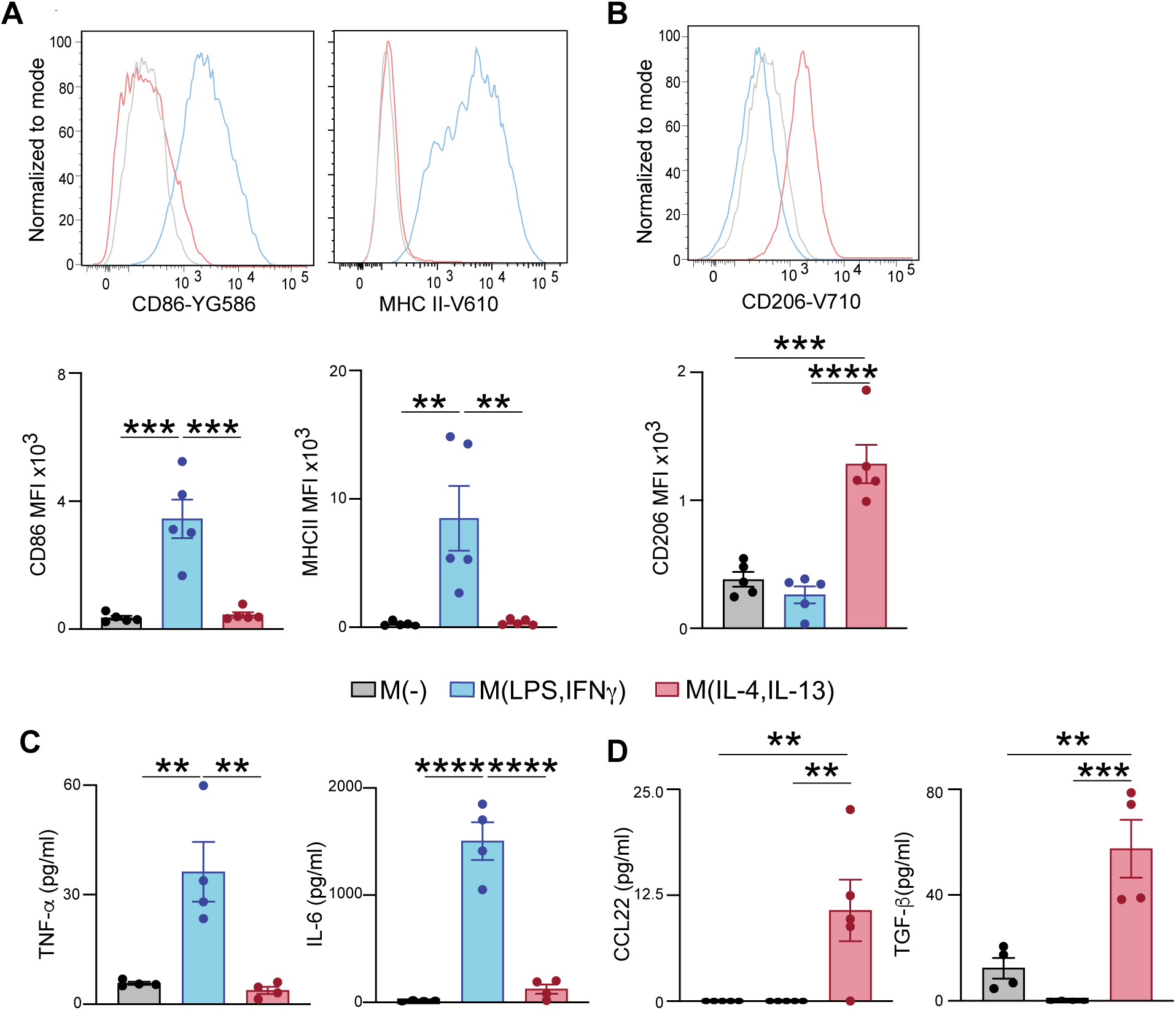
Surface marker expression and cytokine secretion profiles of M(-), M(LPS, IFN-γ), and M(IL-4, IL-13) macrophages. Mouse bone marrow-derived macrophages were differentiated into M(LPS, IFN-γ) or M(IL-4, IL-13) macrophages after incubation with stimulants for 72 hours. Representative and averaged data of **(A)** CD86 and MHC II and **(B)** CD206. Gated on live CD14+CD11b+ cells (n=5). (**C-D)** For cytokine expression analysis, cells were washed with media after polarization, incubated for 24 hours, and supernatants were collected. Amounts of **(C)** TNF-α and IL-6 and **(D)** CCL22 and TGF-β, (n=4 and 5). Bars indicate mean±SEM. Statistical significance was calculated by one-way ANOVA. **p<0.01 ***p<0.001 ****p<0.0001

We also measured the secretion of cytokines, finding that, as expected [30] [32], M(LPS, IFN-γ) macrophages secreted more TNF-α and IL-6 compared to M(IL-4, IL13) macrophages **[Fig. 1C]** [30], which upregulated the expression of anti-inflammatory CCL22 and TGF-β **[Fig. 1D]** [32]. The low expression of inflammatory markers and cytokines, coupled with the high expression of anti-inflammatory factors in M(IL-4, IL13) macrophages, confirmed that these cells were a suitable model of regenerative macrophages.

We next co-cultured BMDMs of the M(-), M(LPS, IFN-γ), and M(IL-4, IL-13) subtypes with hand-picked, intact islets for 6 days. The extended culture time was chosen to allow islet cell death to occur, enabling observation of the potential protective effect of macrophages. We found that islets co-cultured with M(IL-4, IL-13) macrophages had a significantly higher proportion of viable (CytoCalcein^+^Apopxin^-^) cells, and decreased apoptotic (Apopxin^+^) cells, compared to islets cultured without macrophages **[Fig. 2A, S1B]**. By contrast, co-culture of M(-) and M(LPS, IFN-γ) macrophages with islets did not have significant effects on islet cell viability or apoptosis **[Fig. 2A, S1B]**. Similar results were found if islets were dispersed with Accutase to single cells prior to co-culturing with macrophages, with a significant benefit of M(IL-4, IL-13) macrophages on islet cell survival **[Fig. S2, S1B]**.

**Figure 2.**
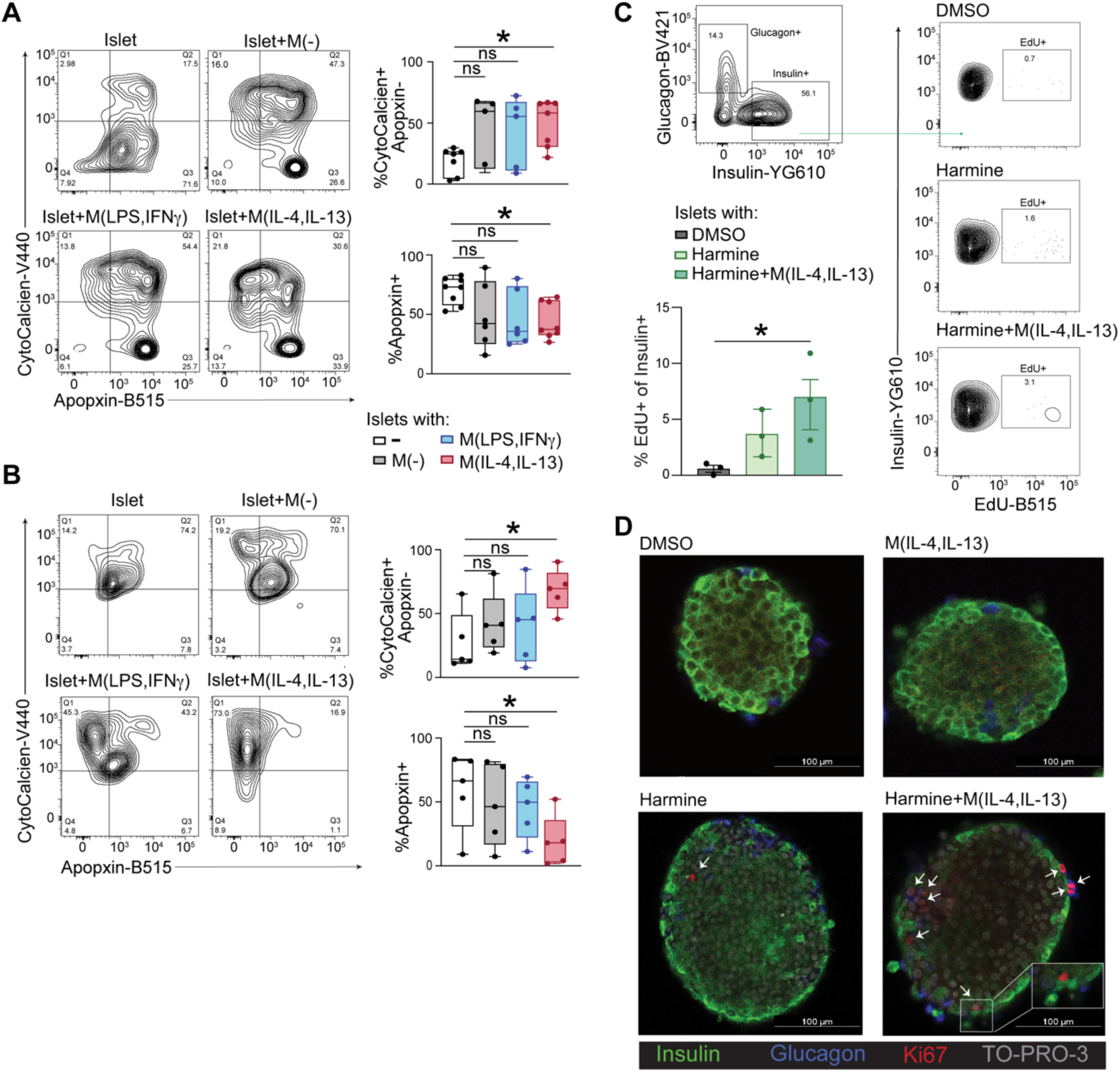
Co-culture with macrophages enhances mouse islet survival. **(A)** Intact mouse islets were directly co-cultured with M(-), M(LPS, IFNγ), and M(IL-4, IL-13) macrophages for 6 days. Representative and averaged data showing the proportions of live (CytoCalcien+Apopxin-) or apoptotic (Apopxin+) islet cells following direct co-culture with mouse macrophages. Gated on CD45-islet cells (n=4-8). **(B)** Intact islets were indirectly co-cultured with macrophages using a transwell system for 6 days. The percentage of live (CytoCalcien+ Apopxin-) mouse islet cells and the percentage of apoptotic (Apopxin+) mouse islet cells on gated on CD45-islet cells (n=5). **(C)** Intact islets were directly co-cultured with macrophages for 3 days. Insulin+ cells were gated as CD45-Glucagon-cells. Representative data showing amount of EdU incorporation into insulin+ beta cells after 3 days of incubation with harmine (10μM, dissolved in 0.1% DMSO) and M(IL-4, IL-13) macrophages (n=3). EdU was added at 10 μM during the final 24 hours. **D)** Immunofluorescence staining of intact mouse islets directly co-cultured with macrophages for 3 days showing the expression of insulin (green), glucagon (blue), Ki67 (red), and TO-PRO-3 (gray), a fluorescent DNA stain, with or without incubation with harmine and M(IL-4, IL-13) macrophages. Statistical significance was calculated by one-way ANOVA. *P value <0.05.

To investigate whether the effect of M(IL-4, IL-13) macrophages on islet cell survival was contact-dependent, we established an indirect islet-macrophage co-culture system using Transwell plates. Similar to observations in direct co-cultures, islets cultured adjacent to M(IL-4, IL-13) macrophages in Transwells exhibited an increased proportions of viable cells and decreased proportions of apoptotic cells **[Fig. 2B, S1B]**. Indirect co-culture with M(-) and M(LPS, IFN-γ) macrophages had no effect on the proportions of Apopxin^+^ or CytoCalcein^+^ islets **[Fig. 2B, S1B]**. Thus, the protective effect of M(IL-4, IL13) macrophages on islet cell survival does not require contact, demonstrating that the observed pro-survival effect is not simply due to phagocytosis of dying/dead islets and rather is likely related to macrophage-derived secreted factors.

We next examined whether M(IL-4, IL-13) macrophages could promote beta cell proliferation. We found that direct co-culture of M(IL-4, IL-13) macrophages with intact islets for 72 hours caused a small but insignificant increase in beta replication, as measured by EdU incorporation **[Fig. S3, S1C]**. Recent studies have shown that small molecules inhibiting dual tyrosine-regulated kinase 1A (DYRK1A), such as harmine and similar compounds, can induce human beta cell replication with rates of 0.25-2.5% [33–35]. We therefore also tested if M(IL-4, IL-13) macrophage effects could be enhanced by harmine. We found that harmine slightly increased the proportion of EdU-positive beta cells, but the effect was not statistically significant. However when in combination with M(IL-4, IL-13) macrophages, there was a synergistic effect, evidenced by significantly higher beta cell replication compared to islets with 0.1% DMSO as vehicle control **[Fig. 2C, S1C]**. The EdU incorporation data were supported by immunofluorescence staining of islets cultured in the presence of harmine and M(IL-4, IL-13) macrophages, which showed an increase in Ki-67^+^ insulin^+^ cells compared to both vehicle control or islets treated with harmine alone **[Fig. 2D]**. Overall, these data suggesting that DYRK1A inhibition can unmask the pro-proliferative effect of M(IL-4, IL-13) macrophages on beta cells.

### Mouse M(IL-4, IL13) macrophages improve beta cell C-peptide secretion

We next asked whether M(IL-4, IL-13) macrophages affect beta cell function. Intact, hand-picked mouse islets were co-cultured with different subtypes of macrophages for 72 hours, then glucose- and KCl-stimulated C-peptide secretion were measured by ELISA. We found that compared to islets alone, or those cultured with M(-)- or M(LPS, IFN-γ) macrophages, islets co-cultured with M(IL-4, IL-13) macrophages resulted in increased C-peptide secretion in response to 16.7 mM glucose **[Fig. 3A]** and KCl **[Fig. 3B]**, with no effect on total islet C-peptide content **[Fig. 3C]**. Similar results were observed when macrophages and islets were co-cultured for 72 hours using a Transwell system **[Fig. 3D-F]**, indicating that cell-cell contact is not essential for the beneficial effect of M(IL-4, IL-13) macrophages on islet function. Since no change in total C-peptide content was observed following direct or indirect co-culture of islets with M(IL-4, IL13) macrophages, their effect on islet function likely involves the modulation of pathways related to insulin secretion and not synthesis.

**Figure 3.**
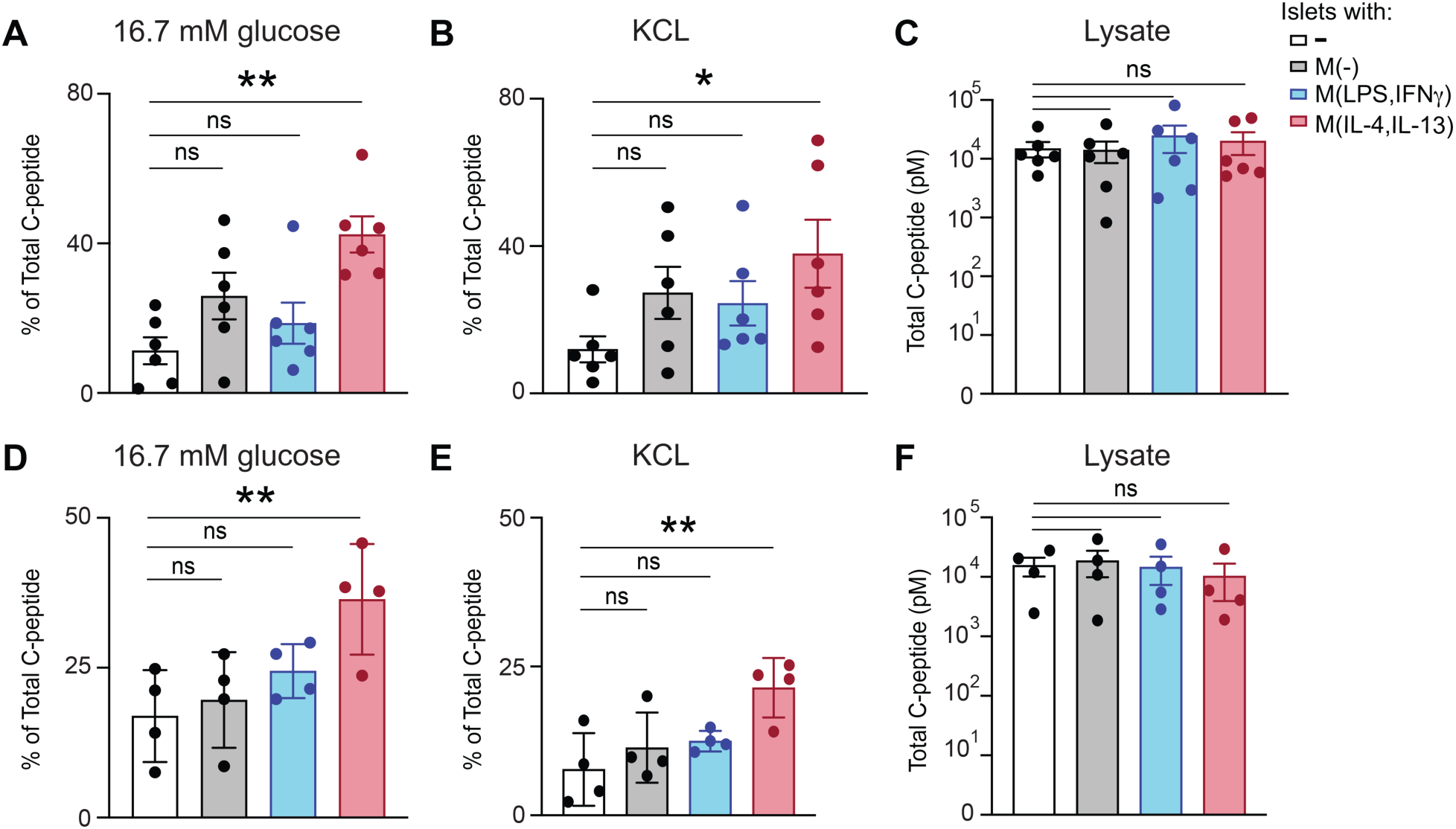
Mouse M(IL-4, IL-13) macrophages enhance islet secretion of C-peptide. Intact mouse islets were co-cultured with M(-), M(LPS, IFNψ), and M(IL-4, IL-13) macrophages for 3 days then were subjects to a glucose-stimulated insulin secretion (GSIS) assay. **(A-C)** C-peptide released into the supernatants was measured by ELISA after stimulation with **(A)** 16.7 mM glucose and **(B)** KCl, and expressed as a percentage of total islet C-peptide content. **C)** Total C-peptide content was measured from islet lysates at the end of the assay (n=6). **(D-F)** Similar measurements were performed using a Transwell co-culture system, where mouse islets were indirectly cultured with macrophages for 3 days and then stimulated with **(D)** 16.7mM glucose then **(E)** KCL. (**F)** Total islet C-peptide content was measured from lysates at the end of the assay (n=4). Bars indicate mean±SEM. Statistical significance was calculated by one-way ANOVA. *p<0.05 **p<0.01

### M(LPS, IFN-γ) and M(IL-4, IL-13) macrophages upregulate markers of regeneration during islet co-culture

Due to their plasticity, macrophages adopt different phenotypes depending on their microenvironment [5, 36]. We hypothesized that, upon co-culturing with islets, macrophages may further shift toward a regenerative phenotype. As expected, after 72 hours of direct co-culture with islets, M(IL-4, IL-13) macrophages showed enhanced expression of the anti-inflammatory marker, CD206 **[Fig. 4A]** along with CCL22 [**Fig. 4B]**, a chemokine produced by regenerative macrophages that regulates immune responses and controls inflammation [37–39]. We found the expression of CD206 and CCL22 was also elevated in inflammatory M(LPS, IFN-γ) macrophages after 72 hours of direct co-culture with islets, suggesting a shift toward a regenerative phenotype **[Fig. 4A&B]**. However, M(-) macrophages did not show upregulation of anti-inflammatory genes, suggesting that the shift toward the regenerative phenotype is specific to polarized macrophages **[Fig. 4A&B]**. Expression of pro-inflammatory cytokines, such as TNF-α, was unaffected by exposure of macrophages of any phenotype to islets **[Fig. 4C]**. Indirect co-culture in the Transwell system did not lead to enhanced expression of CD206 **[Fig. 4D]** or TNF-α **[Fig. 4F]** in either M(LPS, IFN-γ) or M(IL-4, IL-13) macrophages, but did lead to a slight, but insignificant, upregulation of CCL22 expression **[Fig. 4E]**, suggesting that contact dependent mechanisms might regulate the enhanced regenerative phenotype.

**Figure 4.**
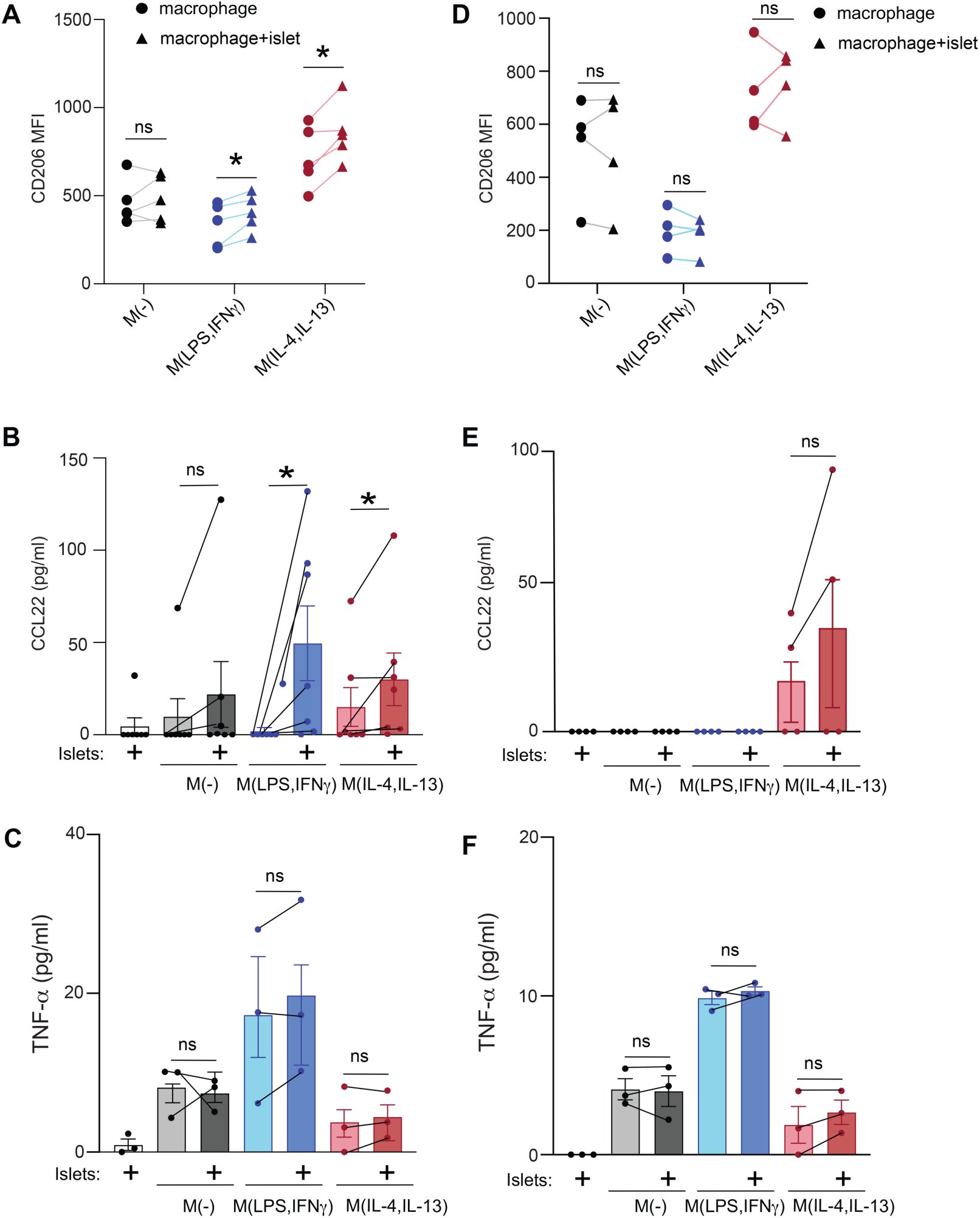
Culturing M(LPS, IFNψ) and M(IL-4, IL-13) mouse macrophages directly with islets increases CD206 expression and secretion of CCL22. **(A-C)** M(-), M(LPS, IFNγ), and M(IL-4, IL-13) macrophages were directly cultured with or without intact islets for 3 days. Live CD14+CD11b+ cells were analyzed for **(A)** CD206 expression (n=5), or supernatants were assessed for secretion of (**B)** CCL22 (n=7) or (**C)** TNF-α (n=3). (**D-F)** M(-), M(LPS, IFNγ), and M(IL-4, IL-13) mouse macrophages were indirectly cultured with or without intact islets using a transwell system for 3 days. **(D**) CD206 expression was measured on live cells gated as CD14+CD11b+ (n=4). Supernatants were analyzed for section of (**E)** CCL22 (n=4) or (**F)** TNF-α (n=3). Bars indicate mean±SEM. Statistical significance was calculated using a one-way ANOVA for A and D, and multiple Wilcoxon test for B, C, E, and F. *p<0.05.

### Human M(IL-4, IL13) macrophages promote human beta cell survival and function

To assess whether macrophage co-culture also impact human beta cell survival and function, we next generated human M(IL-4, IL-13) macrophages by exposure of M(-) macrophages to IL-4 and IL-13 for 72 hours. The resulting cells exhibited high expression of CD206 and CCL17, a homolog of CCL22 **[Fig. 5A&B, S4A]**, confirming successful generation of human macrophages with a M(IL-4, IL13) phenotype. Human M(IL-4, IL13) macrophages were then cultured with hand-picked human islets for 72 hours, and similar to results with mouse cells, we found the proportion of viable (insulin^+^, FVD^-^) beta cells was significantly higher **[Fig. 5C, S4B]**. Upon examination of proliferation, we found that culture with M(IL-4, IL13) macrophages alone did not change the proportion of proliferating (insulin^+^EdU^+^) human beta cells **[Fig. 5D, S4C]**. However, consistent with data from mouse cells, addition of harmine to cultures of human islets and M(IL-4, IL13) macrophages resulted in a small, but significant increase in EdU incorporation **[Fig. 5D, S4C]**. Human islets also exhibited higher glucose- and KCl-stimulated C-peptide release following co-culture with M(IL-4, IL-13) macrophages, without affecting islet C-peptide content **[Fig. 5E-G]**. Consistent with our findings in mouse cells, human M(IL-4, IL-13) macrophages promote β cell survival, function, and proliferation in the presence of harmine.

**Figure 5.**
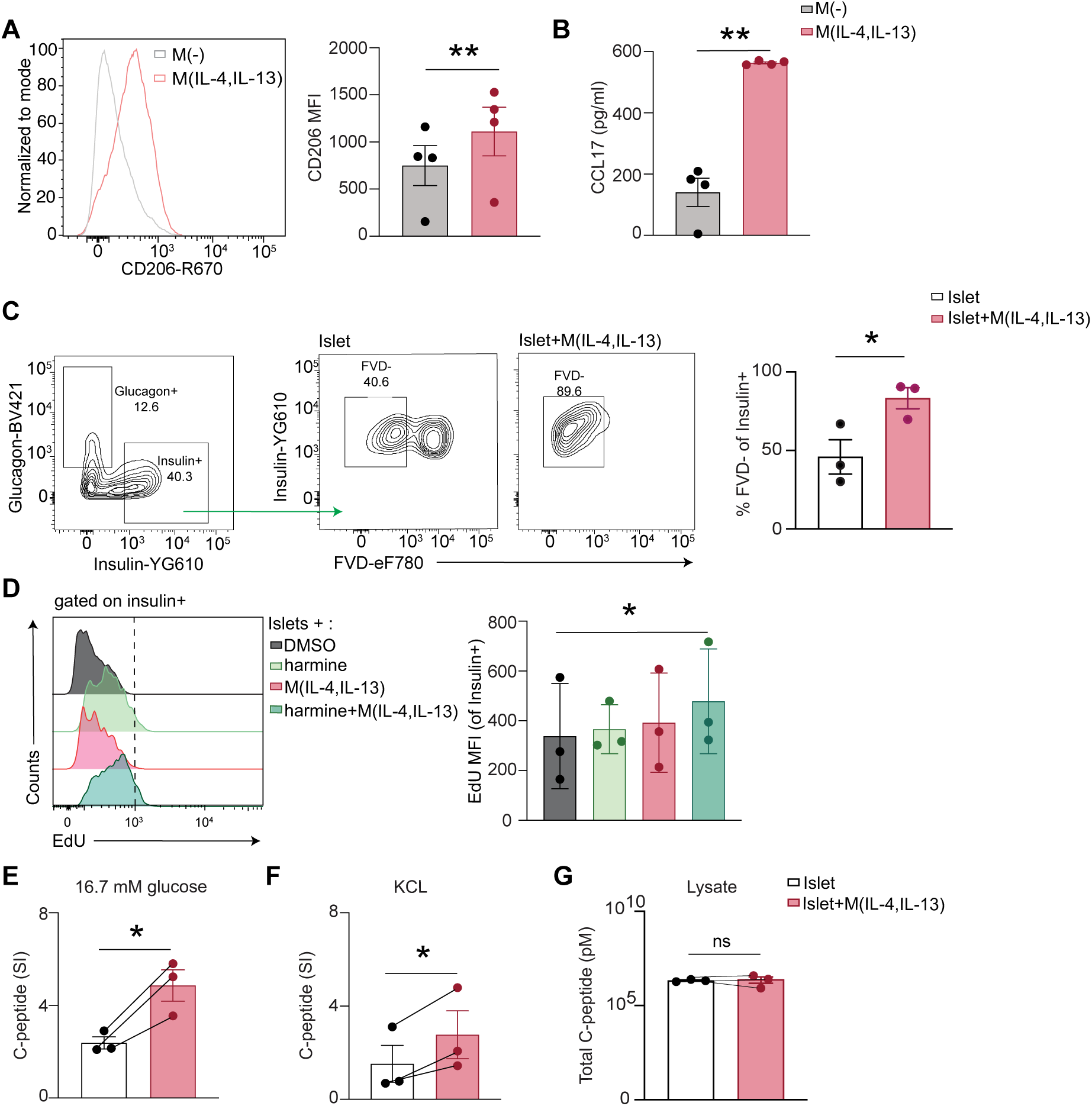
Human M(IL-4, IL-13) enhance human beta cell survival and function. **(A-B)** Human monocyte-derived macrophages were left unpolarized or differentiated into M(IL-4, IL-13) macrophages. (**A)** Representative and averaged data showing CD206 expression on gated live CD14+CD11b+ cells. **(B)** Polarized cells were washed with media, incubated for 24 hours, and then supernatants were collected to measure CCL17 (n=4). **(C-D)** Intact human islets were cultured alone or directly co-cultured with M(IL-4, IL-13) macrophages for 3 days. **(C)** Insulin+ cells were gated on CD45-Glucagon-cells, and the proportion of live (fixable viability dye [FVD] negative) Insulin+ cells was measured (n=3). (**D)** Representative data showing amount of EdU incorporation into insulin+ beta cells after 3 days of incubation with harmine (10μM, dissolved in 0.1% DMSO) and M(IL-4, IL-13) macrophages (n=3). EdU was added at 10 μM during the final 24 hours. Representative and averaged data showing EdU+ expression within insulin+ cells, gated as live CD45-Glucagon- (n=3). (**E-G)** C-peptide secretion by islets cultured with or without M(IL-4, IL-13) macrophages for 3 days. Islets were pre-incubated in low glucose (1.67 mM) before stimulation with (**E)** high glucose (16.7 mM) then **(F)** KCL. (**G)** The level of total C-peptide content in the islets was measured in lysates made at the end of the assay (n=3). The stimulation index (SI) was calculated as C-peptide secretion during high glucose and KCL incubations relative to the initial low glucose condition (n=3). Bars indicate mean±SEM. Statistical significance was calculated by paired t-test for **(A-C)** and **(E-G)** or Friedman test for **(D)**. *p<0.05 **p<0.01

## Discussion

In this study, we tested the ability of macrophages to enhance islet cell survival, regeneration, and function. We found that co-culture of mouse or human islets with macrophages that had been previously skewed to a regenerative phenotype led to improved beta cell survival and function, as well as increased beta cell proliferation in the presence of harmine. Other macrophage phenotypes had more modest, insignificant effects on islet viability and function. These data support the notion that pro-regenerative macrophages can have strong protective and proliferative effects on beta cells.

While co-culture of islets with M(IL-4, IL-13) macrophages or treatment with harmine alone showed a trend toward increased beta cell proliferation, a significant effect in both mouse and human beta cell proliferation was observed only when the two were combined. Harmine belongs to the class of dual-specificity tyrosine phosphorylation-regulated kinase 1A (DYRK1A) inhibitors, which have been shown to induce β cell proliferation at low doses [40]. DYRK1A inhibitors promote β cell proliferation by inducing the nuclear translocation of NFAT transcription factors, which subsequently activate genes that drive cell cycle and suppress those that inhibit it. [40]. More recent findings suggest that combining a DYRK1A inhibitor with either a TGF-β superfamily inhibitor or a GLP-1 receptor agonist further enhances β cell replication [34] [41].

Our data provide a new approach to enhance the effects of DYRK1A inhibition by combining with M(IL-4, IL-13) macrophages, which likely have multiple beneficial properties. Specifically, macrophages are inherently adaptable to their environment, can be engineered to home to specific tissues, act through multiple molecular pathways, and, as cells, may provide more durable therapeutic effects. While the exact mechanisms by which M(IL-4, IL-13) macrophages enhance beta cell proliferation remain to be defined, it is possible that they activate specific signaling pathways in beta cells upon DYRK1A inhibition, similar to GLP-1 agonists. One known mechanism of beta cell proliferation mediated by synergy between GLP-1 and harmine is through activation of cAMP signaling [41]. Interestingly, a previous study has shown that in the gastrointestinal tract, M(IL-4, IL-13) macrophages activate cAMP signaling in epithelial cells following mucosal injury, leading to their proliferation and wound repair [42]. Taken together, our data suggest that M(IL-4, IL-13) macrophages promote beta cell proliferation, and that this effect is revealed by DYRK1A inhibition in both human and mouse islets,. Further studies are needed to elucidate the underlying mechanisms, with the goal of developing novel regenerative therapies for diabetes.

Our findings align with, and expand upon, previous work investigating macrophage-mediated effects on beta cell proliferation [43]. Xia et al. demonstrated that indirect co-culture of mouse primary beta cells with macrophages derived from pancreatic duct-ligated (PDL) mice significantly enhanced beta cell proliferation, even in the absence of DYRK1A inhibitors. However, the PDL model artificially induces local inflammation and promotes islet neogenesis, a process not typically present under normal physiological conditions. Consequently, the macrophages were shaped by an inflammatory microenvironment unique to PDL-induced injury [43]. In contrast, our data reflect the intrinsic capacity of macrophages to promote beta cell proliferation independent of pre-existing pancreatic injury or experimentally induced islet regeneration. Additionally, in our direct co-culture system, physical interaction between macrophages and beta cells may lead to phagocytosis of newly formed or dying beta cells, potentially moderating the overall proliferative response. Together, these differences in experimental models and culture systems likely contribute to the more robust proliferative effect observed in their study.

It is well established that mouse β-cells exhibit a higher proliferative capacity than human β-cells [44–46]. Our data are consistent with this fact since the pro-proliferative effect of M(IL-4, IL-13) macrophages co-culture is higher in mouse than human β-cells. Most mouse studies, including ours, use young animals, expected to have higher beta cell proliferative capacity. In contracts, human islets are typically from older adults and exhibit donor-to-donor variability, influenced by factors such as age, cause of death, and genetic background [47]. Together, these factors should be considered when interpreting cross-species differences in β-cell proliferative responses.

We also sought to evaluate the effect of macrophages on islet cell apoptosis. Islets were cultured with and without macrophages for extended periods of time: 6 days for mouse islets and 3 days for human islets. The shorter culture period for human islets was selected due to the time required for pancreas processing and islet isolation at IsletCore (University of Alberta), as well as the shipping time to our facility. Culture of isolated islets is known to be associated with islet cell death [48]. During isolation, islet cells undergo significant structural and functional changes, including the loss of cell-cell interactions and the absence of trophic factors, which may contribute to reduced islet cell survival [49, 50]. This model offers a useful approach for studying the impact of macrophages on islet survival in the context of type 1 diabetes, allowing us to explore how macrophages interact with islets independently of other external factors. We observed a reduction in islet cell apoptosis and cell loss when islets were co-cultured, both directly and indirectly, with M(IL-4, IL-13) macrophages. These data suggest that factors produced by macrophages, and/or beta cell-macrophage cell-to-cell contact, can not only support beta cell proliferation but can also inhibit beta cell death. While the observed decrease in islet cell death could be partially attributed to phagocytosis of dead or dying cells by neighboring macrophages, the persistence of this effect in the Transwell system suggests it is primarily mediated by soluble factors secreted by macrophages. External factors such as free fatty acids and glucose can induce islet cell apoptosis in vitro [51, 52]. However, in human islet cells, cytokine-induced injury appears to be the primary cause of cell death; for example, IL-1β produced by beta cells can upregulate FAS expression and trigger apoptosis [53]. The mechanism by which macrophages may inhibit islet cell apoptosis, whether by modulating external factors or regulating cytokine responses, remains to be elucidated.

Our findings showing a shift in macrophage phenotype during co-culture with islets to a pro-reparative phenotype are consistent with our previous findings in the streptozotocin and db/db mouse models of diabetes [21]. Apoptotic cells induce a shift toward the regenerative phenotype in macrophages [54, 55]. The process of efferocytosis, where macrophages engulf and clear apoptotic cells, likely plays a central role [54], as does exposure to signals released by apoptotic cells such as TGF-β, which activate pathways promoting the expression of regenerative associated markers and cytokines [55], ultimately aiding tissue repair. Interestingly, we observed a shift in IL-4 and IL-13-derived macrophages toward increased regenerative behavior after co-culturing with islets. These data align with other studies showing that IL-4 or IL-13-polarized macrophages alone cannot induce a tissue-reparative phenotype, and that the presence of apoptotic cells is essential for triggering regeneration mechanisms [56]. Notably, we observed this shift toward a more regenerative phenotype only in M(LPS, IFN-γ) and M(IL-4, IL-13) macrophages, whereas M(-) macrophages retained their original state. This effect may be attributed to prior activation priming macrophages to upregulate receptors that facilitate the transition to a M(IL-4, IL-13) phenotype, a response that is less pronounced in unactivated macrophages [57]. Additionally, we observed a more pronounced shift in macrophage phenotype during direct co-culture with islets compared to the Transwell system, suggesting that physical cell contact, potentially involving efferocytosis, plays a critical role. Whether this shift is driven primarily by direct islet-macrophage interactions or by the engulfment of apoptotic particles remains unclear. Future studies should investigate whether activated macrophages further enhance tissue regeneration after interacting with islets, and the factors that may contribute to this process.

In conclusion, M(IL-4, IL-13) macrophages play a unique and beneficial role in enhancing islet performance, primarily mediated by external factors, distinguishing them from other macrophage types. Our findings, in line with previous studies [21, 24] show that regenerative macrophages promote islet performance. Notably, we found that islets can influence activated macrophages, driving them towards a more regenerative phenotype. To gain a deeper understanding of how macrophages interact with islets to enhance their function, further research is needed using in vivo models, as well as exploring the effects of these cells in combination with other cell types. These insights may have important therapeutic implications for type 1 diabetes. Future research will focus on uncovering the underlying mechanisms driving these effects, as well as developing strategies to reprogram macrophages towards a more regenerative behavior for potential use in autoimmune diseases.

## Acknowledgements

We thank G. Soukhatcheva and M. Komba (Department of Surgery, University of British Columbia, Vancouver, BC, Canada) for their assistance with mouse islet isolation. We also acknowledge the Flow Core Facility at BC Children’s Hospital Research Institute (Vancouver, BC, Canada) for support with flow cytometry. We are grateful to Patrick MacDonald and the team at the University of Alberta IsletCore for providing human islets. We also thank Canadian Blood Services for supplying human buffy coats through their Blood for Research program.

## Data availability

Data supporting the findings of this study are available from the corresponding author upon request.

## Contribution statement

C.B. Verchere and M.K. Levings are the guarantors of this work and take full responsibility for the integrity of the data and the decision to publish. C.B. Verchere, M.K. Levings, and S.Q. Crome contributed to the study design and provided overall supervision and guidance. M. Monajemi and A.S. Craciun performed the experiments. M. Monajemi conducted the data analysis and drafted the manuscript. M. Mojibian contributed to data acquisition by assisting with microscope imaging. Q. Huang generated the human macrophages, and L. Dai contributed to mouse islet isolation. All authors critically revised and approved the final manuscript.

## Abbreviations

BMDM: bone marrow-derived macrophages
DYRK1A: dual tyrosine-regulated kinase 1A
GLP-1R: glucagon-like peptide-1 receptor
KRB: Krebs-Ringer Bicarbonate buffer
PBMCs: Peripheral blood mononuclear cells
PDL: pancreatic duct-ligation
TGFβSF: transforming growth factor–β superfamily

