## Supplemental Data for "Human and mouse regenerative macrophages enhance beta cell survival, function, and proliferation"

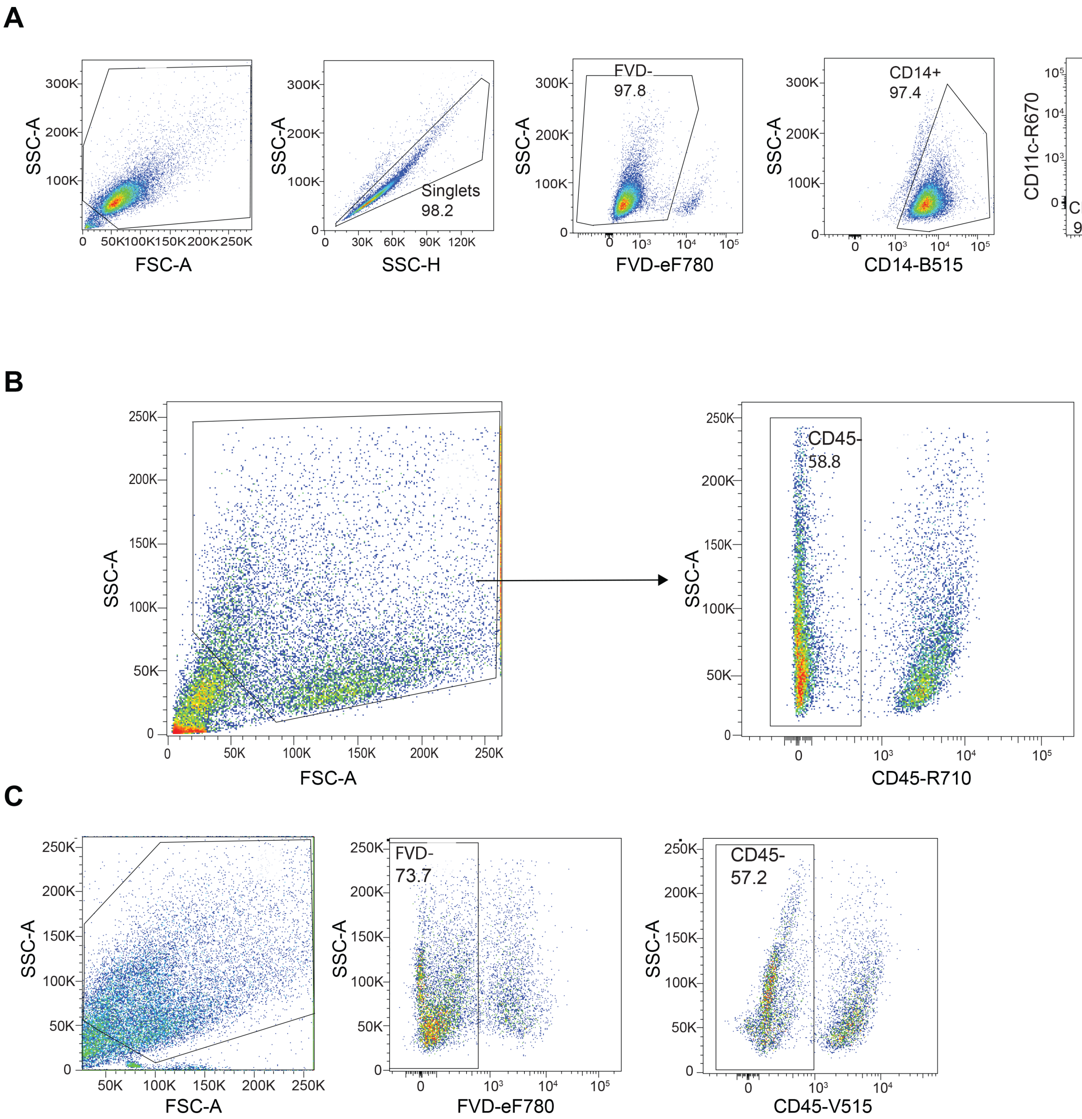


**Supplementary Figure 1. Flow cytometric gating strategies used to identify mouse macrophages and islets. A)** Flow cytometry gating strategy used to pre-gate mouse macrophages which were incubated with or without stimulants (IL-4, IL-13) or (LPS, IFNg) for 72 hours. **B)** Flow cytometry pre-gating strategy used to detect apoptosis in mouse islets cultured alone or co-cultured with M(-), M(LPS, IFNg), and M(IL-4, IL-13) macrophages for 6 days. **C)** Flow cytometric pre-gating strategy used to analyze insulin$^{+}$ cells cultured alone or co-cultured with macrophages for 3 days.

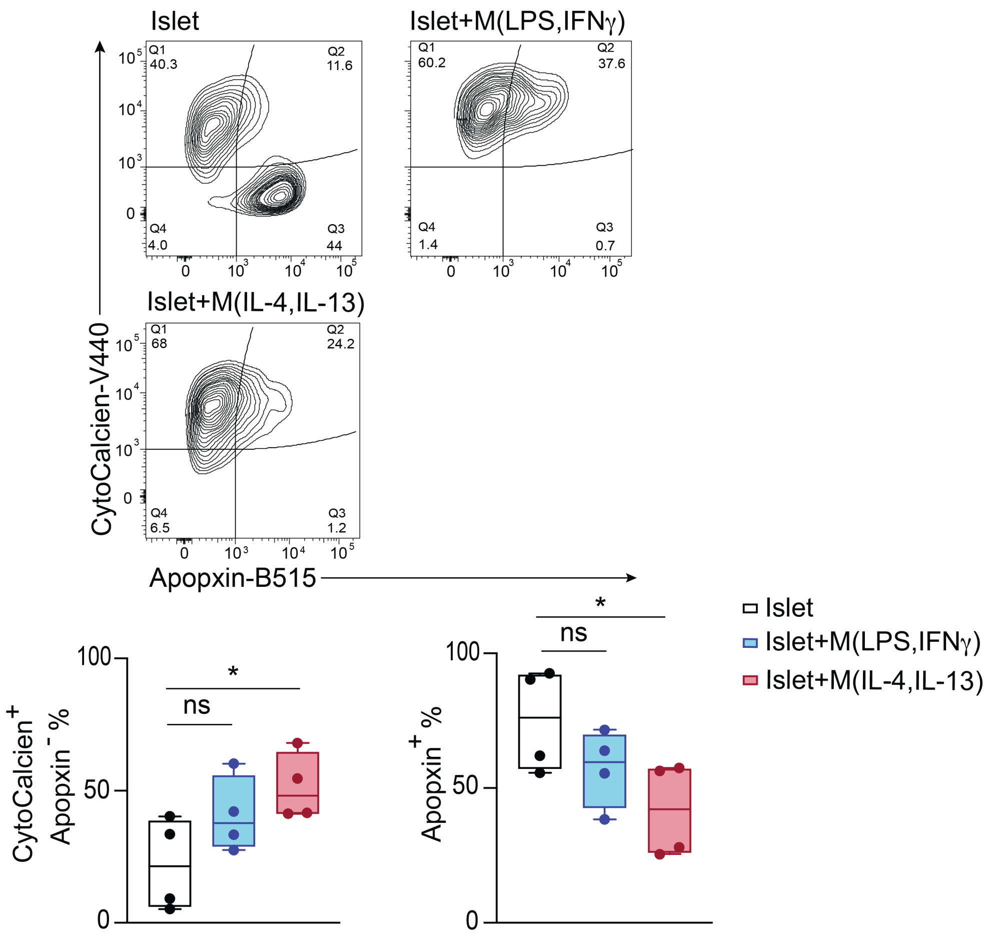


**Supplementary Figure 2. Survival of dispersed islets is enhanced by co-culture with M(IL-4, IL-13) macrophages.** Islets were dispersed using Accutase and directly co-cultured with M(LPS, IFNγ) or M(IL-4, IL-13) macrophages for 6 days. Representative and averaged data showing the proportion of live (CytoCalcien+Apopxin-) or apoptotic (Apopxin+) islet cells gated on CD45-negative cells (n=4). Statistical significance was calculated by One-way ANOVA.*P value <0.05.

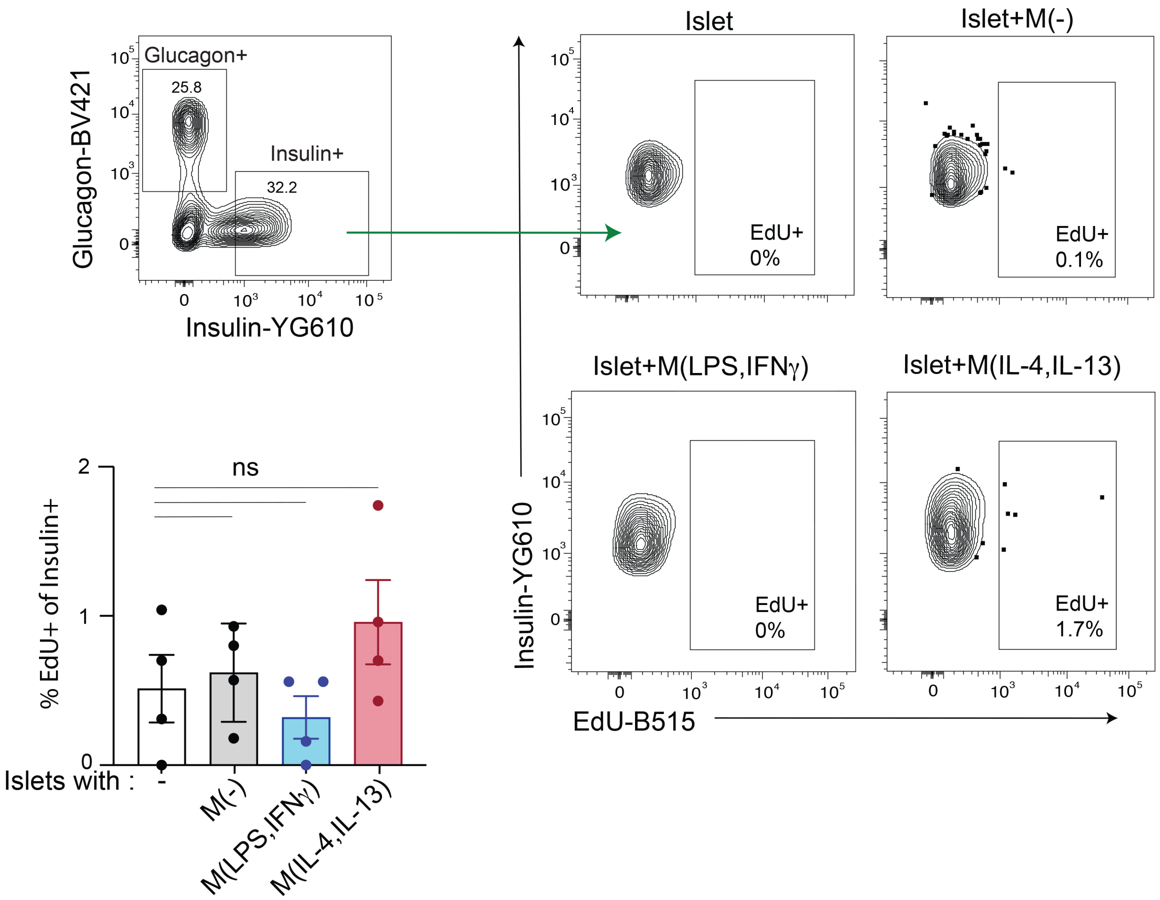


**Supplementary Figure 3. Co-culture with M(IL-4, IL-13) macrophages alone does not significantly promote beta cell proliferation.** Intact mouse islets were directly co-cultured with M(-), M(LPS, IFNγ), and M(IL-4, IL-13) macrophages for 3 days**.** Live insulin+ cells were pre-gated as CD45- cells. Representative data showing proportions of EdU+ cells within insulin$^{+}$glucagon$^{-}$ cells after incubation with the indicated subsets of macrophages (n=4). Bars indicate mean±SEM. Statistical significance was calculated by one-way ANOVA.

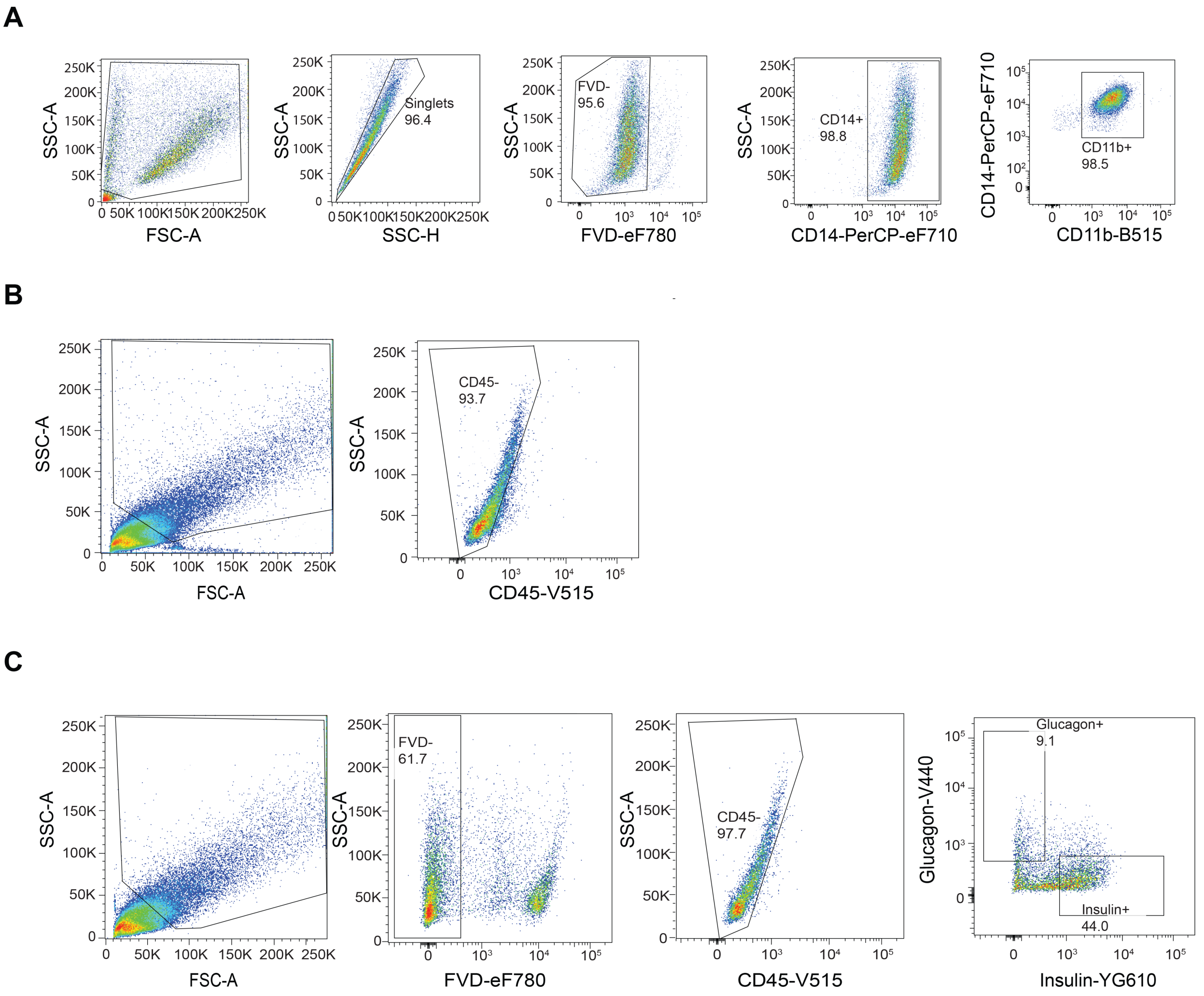


**Supplementary Figure 4. Flow cytometric gating strategies used to identify human macrophages and islet cells. A)** Flow cytometry gating strategy used to identify human macrophage phenotype, incubated with or without stimulants (IL-4, IL-13) for 72 hours. **B)** Flow cytometry pre-gating strategy used to detect apoptosis in human insulin+ cells cultured alone or co-cultured with M(IL-4, IL-13) macrophages for 3 days. **C)** Flow cytometric pre-gating strategy used to identify EdU$^{+}$ insulin$^{+}$ human cells cultured alone or co-cultured with (IL-4, IL-13) macrophages for 3 days.

**Supplementary Table 1. List of antibodies used for flow cytometry in the study.**

| Flow cytometry | Flow cytometry | Flow cytometry | Flow cytometry |
|---|---|---|---|
| CD45 | 30-F11 | AF700 | BioLegend |
| CD45 | 30-F11 | V500 | BD Biosciences |
| CD45 | HI30 | V500 | BD Biosciences |
| CD14 | rmC5-3 | FITC | BD Biosciences |
| CD14 | 61D3 | PerCP-eFluor 710 | BioLegend |
| CD11b | M1/70 | BB700 | BD Biosciences |
| CD11b | CBRM1/5 | FITC | Invitrogen |
| CD11c | HL3 | APC | BD Biosciences |
| CD86 | B7-2 | PE | BD Biosciences |
| MHC Class II | M5/114.15.2 | BV605 | BioLegend |
| CD206 | C068C2 | BV711 | BioLegend |
| CD206 | 19.2 | APC | BD Biosciences |
| Insulin | 182410 | PE-Dazzle | R&D Systems |
| Glucagon | U16-850 | BV421 | BD Biosciences |

**Supplementary Table 2. List of antibodies used for immunofluorescence in the study.**

| Flow cytometry | Flow cytometry | Flow cytometry | Flow cytometry |
|---|---|---|---|
| Glucagon | U16-850 | BV421 | BD Biosciences |
| Insulin | 182410 | PE-Dazzle | R&D Systems |
| Ki67 | SolA15 | FITC | eBioscience |

**Supplementary Table 3. List of human islet deceased donors used in the study.**

| Flow cytometry | Flow cytometry | Flow cytometry | Flow cytometry |
|---|---|---|---|
| R528 | Human islet | Male | 53 |
| R537 | Human islet | Female | 18 |
| R543 | Human islet | Female | 54 |
| R547 | Human islet | Female | 62 |
| R555 | Human islet | Male | 35 |
| R563 | Human islet | Male | 59 |
| R567 | Human islet | Female | 20 |
